# gGN: learning to represent graph nodes as low-rank Gaussian distributions

**DOI:** 10.1101/2022.11.15.516704

**Authors:** Alejandro A. Edera, Georgina Stegmayer, Diego H. Milone

**Affiliations:** Research Institute for Signals, Systems and Computational Intelligence, sinc(i), FICH-UNL/CONICET, Ciudad Universitaria, Santa Fe, Colectora Ruta Nacional No 168 km. 0, Paraje El Pozo, Santa Fe, 3000, Argentina

## Abstract

Unsupervised learning of node representations from knowledge graphs is critical for numerous downstream tasks, ranging from large-scale graph analysis to measuring semantic similarity between nodes. This study presents gGN as a novel representation that defines graph nodes as Gaussian distributions. Unlike existing representations that approximate such distributions using diagonal covariance matrices, our proposal approximates them using low-rank perturbations. We demonstrate that this low-rank approximation is more expressive and better suited to represent complex asymmetric relations between nodes. In addition, we provide a computationally affordable algorithm for learning the low-rank representations in an unsupervised fashion. This learning algorithm uses a novel loss function based on the reverse Kullback-Leibler divergence and two ranking metrics whose joint minimization results in node representations that preserve not only node depths but also local and global asymmetric relationships between nodes. We assessed the representation power of the low-rank approximation with an in-depth systematic empirical study. The results show that our proposal was significantly better than the diagonal approximation for preserving graph structures. Moreover, gGN also outperformed 17 methods on the downstream task of measuring semantic similarity between graph nodes.

## 1 Introduction

To represent facts about the world, knowledge bases use triplets in which a fact is defined as a well-defined relationship between two entities. For example, WordNet [1], Freebase [2], and Wikidata [3] are knowledge bases widely exploited in diverse applications [4, 5, 6]. Another important example is the Gene Ontology (GO) [7, 8] that is widely used for studies aimed at elucidating the diverse roles that genes play in cell biology [9, 10]. Knowledge bases are frequently represented as graphs, where triplets are labeled directed edges between nodes. To exploit such graphs effectively, recent efforts have proposed to use neural networks to represent graph nodes as point vector representations (embeddings) that preserve graph features as much as possible in a lowdimensional space [11]. However, because the underlying graph is generally assumed to be undirected, the learned representations are unable to properly preserve asymmetric relationships in directed graphs [12, 13, 14] and also struggle to model hierarchical structures [15, 16, 17, 18]. At the end, this leads to suboptimal node representations, negatively impacting in downstream tasks.

This study proposes gGN as a novel representation for graph nodes that uses Gaussian distributions to map nodes not only to point vectors (means) but also to ellipsoidal regions (covariances). In contrast to point vectors, the use of distributions enables expressing asymmetric relationships more naturally [14]. Being inspired by previous works [19, 16], the proposed approach built upon them to make three novel contributions. First, the proposed Gaussian distributions are not parameterized by classical diagonal matrices but rather by low-rank covariance matrices, so far underexplored in existing works proposing word or node representations. Unlike diagonal matrices, low-rank covariance matrices have much more flexibility to express dependencies between embedding dimensions [20, 21], enabling the representation of more complex graphs. Although non-diagonal covariance matrices generally scale quadratically with the number of nodes, the low-rank form has a tractable computational cost [22]. Second, we propose a novel loss function based on the reverse Kullback-Leibler (KL) divergence. Besides the KL is well suited for capturing asymmetric local structures, the reverse KL additionally leads to Gaussian distributions whose entropies properly preserve the information contents of nodes. Such preservation not only provides a strong link with classic studies on measuring word similarity based on information theory [23, 24], but also enables us to measure the semantic similarities between nodes in a novel way through the divergences of their corresponding representations. Third, to capture asymmetric global structures, the loss function also incorporates two ranking-based components aimed to asymmetrically preserve the distances between nodes, given by their shortest path lengths. We made the code of gGN publicly available (https://github.com/aedera/ggn) as an easily installable Python package, which can be used for learning node embeddings from scratch.

## 2 Related work

Many different unsupervised approaches have been proposed for learning representations of graph nodes [25, 26, 11]. They are generally divided into two main categories: matrix factorization and random-walk based approaches. Matrix factorization-based methods, such as GraRep [27], construct a high-order proximity matrix that is factorized to obtain low-dimensional node embeddings. A limitation of these methods is that they are not easy to scale up for large graphs. In contrast, this is not a limitation for random walk-based methods, such as DeepWalk [28], LINE [29], node2vec [30] and VERSE [31], which describe the neighborhood of each node as random walk paths that are jointly preserved by a point-vector embedding using an objective like skip-gram [32].

Contrasting with the aforementioned approaches, which represent nodes as point vectors, there are a few methods that are able to represent nodes as probability distributions. One of the first attempts is word2gauss [19] that uses Gaussian distributions as representations. Since this neural network was specifically designed for words, representations are optimized such that the divergence between them preserves word co-occurrences. Once learned, the resulting mean vectors represent the semantics of words, while the covariance matrices describe the uncertainty of meanings. Interestingly, these Gaussian embeddings can very naturally encode asymmetric relationships between words through embedding encapsulation patterns [14], which can effectively express semantic orderings between meanings (e.g. mammal ≺ *Homo sapiens*). Recent works have extended these Gaussian embeddings with a Bayesian strategy to perform automatic word sense disambiguation [33], and with linear combinations of Gaussian distributions to represent subwords [34]. Interestingly, word2gauss has been also extended to general graphs in an approach known as Graph2Gauss [16], where embeddings representing graph nodes are learned by employing unsupervised and supervised strategies. This approach has also shown to be effective as an explanatory tool for analyzing complex real-world graphs [35]. Notably, all these approaches based on Gaussian embeddings have in common that covariance matrices are diagonal, hampering the correct representation of some structures.

## 3 Low-rank Gaussian embeddings

### 3.1 Representation

Given an unweighted directed graph with *n* nodes, the aim is to map every node *i* to a Gaussian distribution 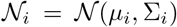, where 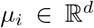 is the mean with *d* dimensions and 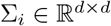 is the covariance matrix. To make computations affordable, the covariance matrix is commonly approximated as a diagonal matrix Σ_*i*_ = *D_i_*. While attractive for its simplicity, this diagonal assumption comes at the cost of overly limiting the range of density forms, preventing complex hierarchical structures from being modeled properly. To overcome these limitations and to preserve computational tractability, this study proposes to approximate the covariance matrix with a low-rank perturbation

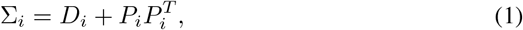

where the diagonal covariance matrix *D_i_* for the node i is perturbed with a r-rank covariance factor 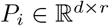. The outer product of the covariance factor with its transpose adds off-diagonal entries to the diagonal matrix. Interestingly, such a perturbation can naturally express richer hierarchical structures through correlations between the embedding dimensions, enabling Gaussian distributions to rotate their densities. Although the cost of the low-rank approximation is higher than the diagonal approximation, it is substantially lower than its full-rank counterpart if *r* ≪ *d*.

To learn these low-rank Gaussian distributions, we propose a neural network consisting of a hidden layer in 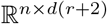. Given as input a one-hot vector in {0,1}^*n*^ for node *i*, it is projected into the hidden layer to obtain 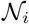. The embedding 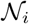 contains the flattened parameters of a Gaussian distribution: the first *d* dimensions are the mean vector *μ_i_*, the following *d* dimensions are the diagonal matrix *D_i_*, and the remaining dimensions are the covariance factor *P_i_*.

### 3.2 Loss function

To learn the weights of the aforementioned neural network, we propose a loss function 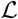 that is calculated from a matrix 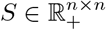. Each entry *S_ij_* is the length of the shortest path from node *i* to node *j*, where *S_ij_* = ∞ if both nodes are unreachable. Given *S* built from an input directed graph, the loss function is locally defined as

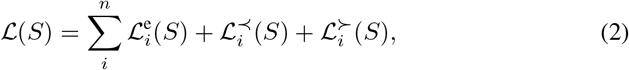

whose components aim to preserve the entailment relationships between nodes 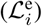, as well as the topology of ancestors 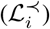 and descendants 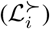.

In this function, the first loss component is

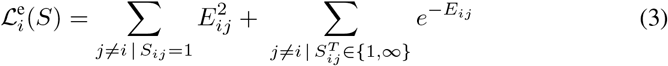

and involves two summations. The left summation is over the parents (*S_ij_* = 1) of node *i*, whereas the right summation is over both its immediate descendants 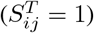 and its remaining unreachable nodes 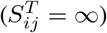. *E_ij_* is a function that assigns a scalar energy [36] to the node pair, which is defined based on the Gaussian distributions 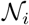 and 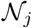 as will be explained in §3.3. Consequently, when minimizing the loss function, this component quadratically reduces the energy between node i and their parents, but exponentially pushes node *i* apart from their immediate descendants and remaining unreachable nodes. This loss component is thus capturing the local structure of a node by preserving its first-order proximities.

The second loss component is defined as a typical mean squared error

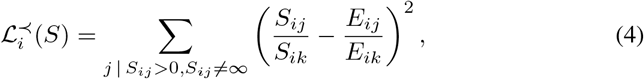

where *k* = argmax_*j*_ *S_ij_*, and *S_ik_* is then the longest shortest path starting from node i. Here, the summation is over the ancestors of *i* (i.e., *S_ij_* > 0, *S_ij_* ≠ ∞). This component assesses how well the energies between nodes *i* and *j* preserve the topology of the ancestors of *i*. This topology is defined by ranking the ancestors according to the lengths of their shortest paths from node i. Note that both shortest path lengths and energies are normalized by using *S_ik_* and *E_ik_*, respectively, to effectively express rankings relative to *i*. This loss component enables embeddings to capture high-order proximity information of each node.

Analogous to the previous one, the last loss component is also a mean squared error

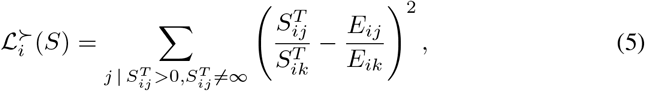

where 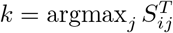, and 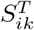 is then the longest shortest path ending at node *i*. The summation here is over the set of descendants of node *i*. Consequently, in contrast to 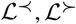 assesses how well the energies between nodes *i* and *j* preserve the topology of the descendants of *i*. This loss component also enables embeddings to capture high-order proximity information asymmetrically with respect to 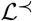.

### 3.3 Energy function

In the loss function, the energy between nodes is concretely defined as the KL divergence between the Gaussian distributions representing these nodes. This divergence has a closed form for such distributions [37]

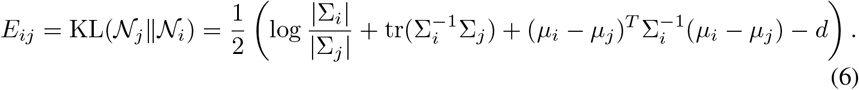

The KL divergence is non-negative and equals zero if both distributions are equal. Although this divergence is not a metric, it generalizes the Pythagoras’ theorem for square distances [38]. KL is high when, for example, there is a region where the density 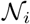 is low but 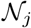 is high. Minimizing the KL will thus promote situations where the density 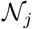 is *encapsulated* within the region where 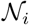 is high, while opposite situations will be penalized. The computation cost of the KL is dominated by the calculation of the determinant and inverse of the *d*-by-*d* covariance matrices. However, it can be largely reduced by exploiting the low-rank form through the use of the matrix determinant lemma [39] and the Woodbury matrix identity [40] (details in §A). Calculations boil down to apply a Cholesky decomposition, with time complexity *O*(r^3^), and then calculating the inverse (forward substitution) and determinant of the resulting Cholesky factor, requiring *O*(*r*^2^) and *O*(*r*) time, respectively.

In contrast to existing approaches [19, 16, 14], this study defines the energy *E_ij_* not as the forward KL, 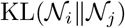, but rather by the reverse KL, 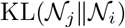. The forward KL leads to Gaussian distributions whose dispersions reflect how abstract the represented concepts are [19, 14]. For example, an abstract concept such as “animal” has high dispersion because it includes multiple more specific concepts with lower dispersions, such as “mammal”, “feline” and “nematode”. In contrast, the reverse KL proposed here has the opposite approach, which is consistent with the perspective of information theory. Since the entropy of a Gaussian distribution is a function of its dispersion, distributions providing more information exhibit high entropy, and vice versa. Interestingly, because the entropy is formally defined as the expected information content of a random variable [41], the higher the entropy, the richer its information content. That is, the deeper a node is, the higher its information content, and thus greater its dispersion and entropy are. Consequently, the use of the reverse KL is in line with previous works where the information content of words is similarly defined [42, 43, 23, 24].

## 4 Experiments

We performed four sets of experiments to evaluate the performance of gGN. The first one studies the node representations that it learned from toy graphs, to clearly illustrate the representational benefits of the low-rank approximation. The second set of experiments assesses the benefits of the low-rank approximation using real-world graphs, and its ability for preserving topologically-relevant graph features. The third set of experiments carries out an ablation study to demonstrate the impact that the three loss components have on obtaining meaningful node representations. The last set of experiments evaluates the performance of low-rank Gaussian embeddings on the important task of measuring the semantic similarity between graph nodes.

### 4.1 A case study on toy graphs

To clearly illustrate the representational benefits of using low-rank Gaussian embeddings, 4 directed graphs were defined to showcase common structural features. They were fed into gGN to learn 2-dimensional Gaussian embeddings of rank 2. Figure 1A depicts a chain graph along with its learned embeddings (top). Here, each (confidence) ellipse depicts the region of a Gaussian embedding/distribution containing points within one standard deviation of the mean. The graph hierarchical relationships are captured through patterns of encapsulation between embeddings such that the densities of parents are within regions in which their children assign high density. Expectedly, the most inner distribution corresponds to the root.

**Figure 1:**
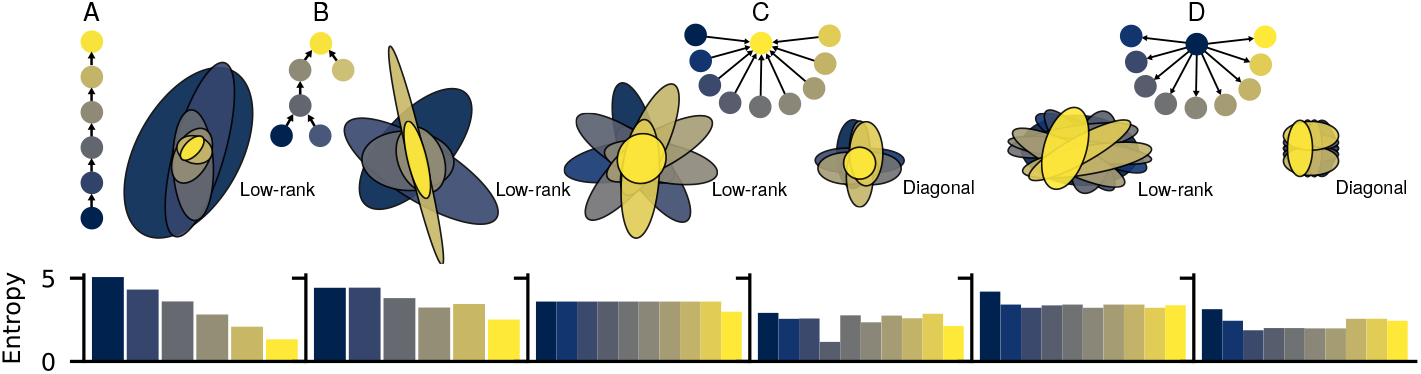
Gaussian embeddings. The nodes of four toy graphs (A-D) are represented through Gaussian embeddings (top). Bars plot the entropy of each embedding (bottom).

To quantify the aforementioned result, Figure 1 also shows the entropies of the embeddings (bottom). Since the entropy of a Gaussian distribution is a function of its covariance matrix, large ellipses are associated with high-entropy distributions, and vice versa. Interestingly, in Figure 1A, since low-entropy distributions correspond to nodes near the root, the entropy is preserving information about node depth. Moreover, because the entropy is defined as the expected information content for a random variable, low-entropy distributions convey low information contents, and thus their nodes can be interpreted as representing more abstract concepts. Notably, Figure 1B shows that similar results are reached when using a more complex graph. Here, encapsulations between embeddings can successfully preserve the two different branches of the new graph.

The role that covariance matrices play can be clearly appreciated in Figure 1C and 1D, where nodes are represented by low-rank (left) and diagonal (right) covariance matrices. In graph C, unlike the diagonal case, the low-rank ellipsoids of the children nodes display varying degrees of rotation. This leads to rather homogeneous entropies among children that are also higher than that of the root. Similar tendencies can be observed from the embeddings in Figure 1D. These results highlight the meaningful role that low-rank covariance matrices have in representing nodes.

### 4.2 Loss convergence for the low-rank approximation

To investigate the learning stability of the low-rank approximation, we analyzed the loss curves yielded by gGN when learning 10-dimensional Gaussian embeddings using spherical, diagonal and low-rank covariance matrices, with rank values ranging from 1 to 4. This analysis used three real-world graphs (DAGs) obtained from the GO [7]: Biological Process (BP), Cellular Component (CC) and Molecular Function (MF). These graphs were selected because their complexity, despite being not trivial, is suited for systematic in-depth analysis with our computational budget. Data and training details are provided in §B.

The obtained loss curves are shown in Figure 2 and, regardless of the graph, a dramatic drop is observed when using the diagonal instead of the spherical approximation, demonstrating the importance of the covariance matrix in model optimization. Similarly, the curves of the low-rank approximations (blue) are significantly better than those of the spherical and diagonal approximations (green). Moreover, the curves also show that the higher the rank is, the lower the loss values are, underlining the benefits that the low-rank covariance matrix has during learning. Interestingly, higher rank values tend to give modest or marginal loss improvements. Since computations are cheaper for lower ranks (see §C), this result highlights that good and stable loss curves can be obtained with a low computational cost.

**Figure 2:**
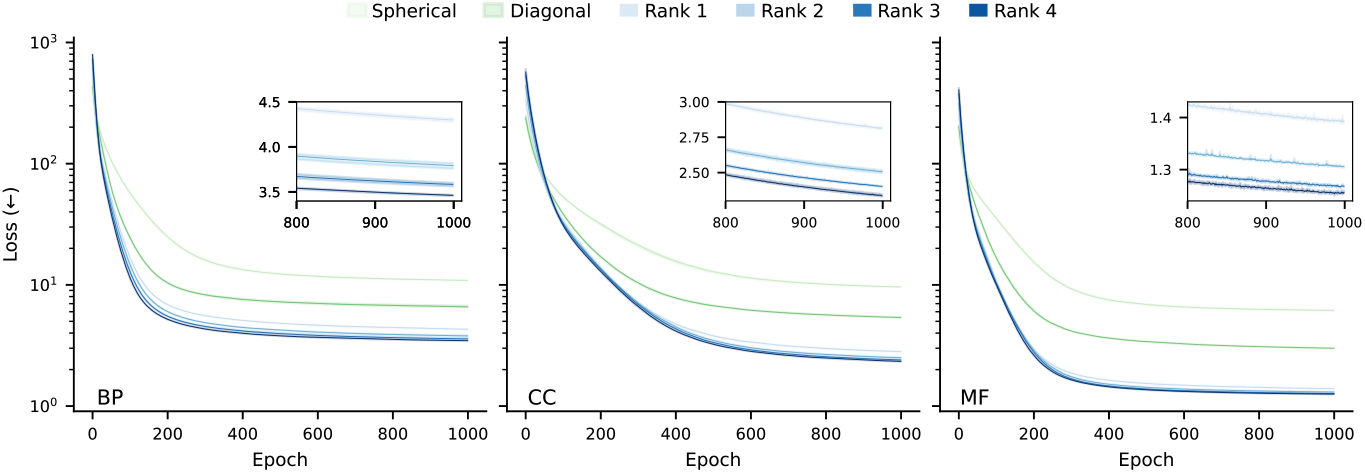
Loss curves for different covariance matrices on three real-world directed graphs. Curves depict the average values and shaded regions show the dispersion for different seeds. Zoom-in figures better visualize differences among ranks.

Since Gaussian distributions showed very stable loss curves at very low ranks, we further investigated whether this tendency was still held for higher embedding dimensionalities. To this aim, we analyzed the loss curves yielded when learning spherical, diagonal and rank-1 approximations on BP, CC and MF by ranging the embedding dimension d from 10 to 50. Figure 3 shows the resulting curves, where the losses of the rank-1 embeddings (blue) are always better than those of the spherical and diagonal ones (green), regardless of the embedding dimensionality. For example, the spherical and diagonal embeddings with the highest number of dimensions (*d* = 50) are completely unable to obtain better losses than all rank-1 embeddings, even than those using the lowest number of dimensions (*d* = 10). On the other hand, among the rank-1 embeddings, small loss gains are observed when dimension *d* is higher than 10, indicating that the embedding dimensionality is not a significant factor. Based on these results, *d* =10 was selected as an experimentally appropriate and computationally affordable embedding dimension for the following experiments.

**Figure 3:**
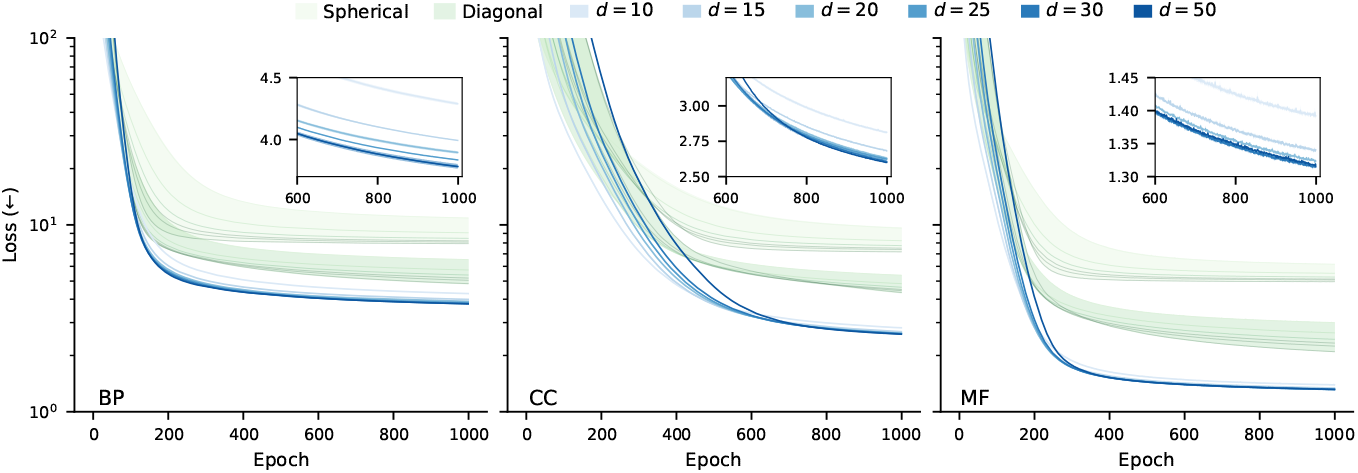
Impact of embedding dimensionality. Loss curves for different embedding dimensions *d*. The green-shaded region depicts curves yielded by spherical and diagonal embeddings, whereas blue curves are rank-1 embeddings.

### 4.3 Preservation of graph features

To assess the quality of the Gaussian embeddings learned by gGN from the real-world graphs, we evaluated whether they were able to preserve two important graph features of nodes: the lengths of the shortest paths between them and their depths. To contextualize this evaluation, we also included the node embeddings learned by Graph2Gauss [16], which is a strong baseline as it uses Gaussian distributions as representations.

First, to assess whether the shortest path lengths were properly preserved by embeddings, we measured how correlated such lengths between nodes were with the KLs between their corresponding Gaussian embeddings. Since the KL is asymmetric, this correlation was also measured after inverting the directions of all graph edges. In the following, we will use ≻ to indicate results with the original graph directions and ≺ to indicate the results with the inverted directions. Correlations were quantified by calculating the Pearson coefficient (shown in Table 1). As expected, all embeddings show a strong positive correlation: the longer the length of the shortest path between two nodes, the higher the KL between their corresponding node representations (detailed plots are provided in §D). More importantly, this positive correlation is held for both directions of the KL (≻ and ≺), indicating that the global structure is asymmetrically preserved. In general, the diagonal Gaussians are statistically similar to or better than the spherical ones, in particular for node depth. However, there is a case where the spherical Gaussians outperform the diagonal ones (for ≻ on BP). This can be expected because, as it will be shown in the ablation study below, preserving node depth information has a higher impact on the loss function than preserving the shortest path lengths, and diagonal models are better suited to preserve the former one. Therefore, when the parameters allow the model to maximize performance on node depth, all the expressiveness of the Gaussian distribution is used for preserving the most important loss component, at the cost of neglecting the other two ones. Notably, on the three graphs the highest correlations are obtained by the low-rank approximations proposed here. In particular, low-rank models showed to be much better than Graph2Gauss on node depth. These results indicate that the asymmetric relationships between nodes, measured by the divergences between their embeddings, are effectively preserving the asymmetric topology of the directed graphs, as defined by the shortest path lengths.

**Table 1:**
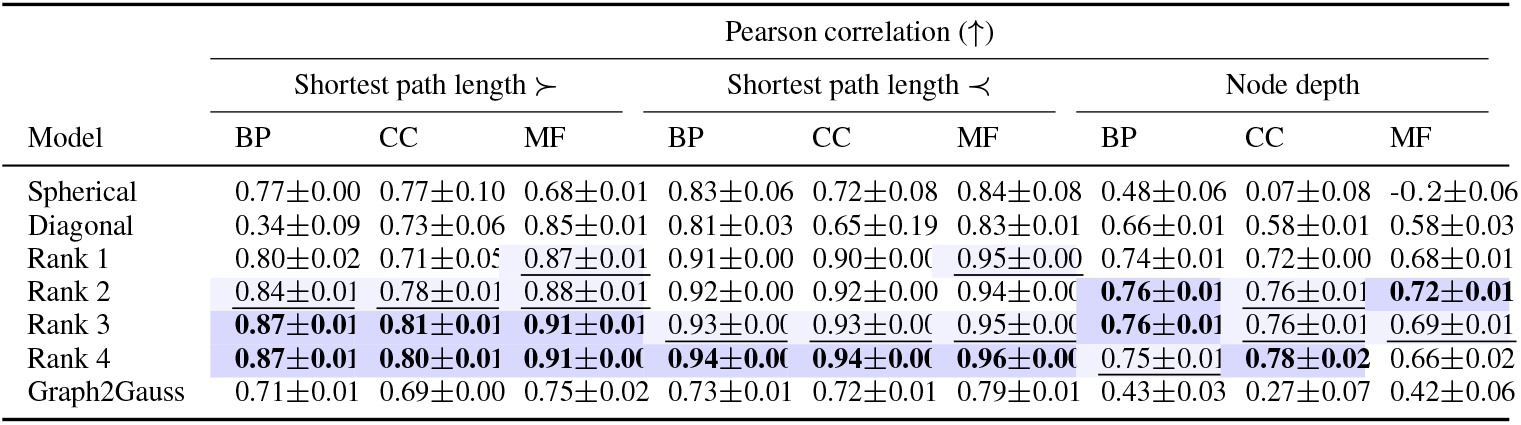
Graph features preserved by Gaussian embeddings. The two best values are first boldfaced and then underlined, respectively.

Second, to assess whether node depths were properly preserved, we measured how well such depths were correlated with the entropies of their corresponding embeddings. Here, the depth of a node was defined as the number of its ancestors. Correlations were also quantified by computing the Pearson coefficient. The results in Table 1 reveal that almost all the embeddings show a positive correlation on the three graphs: the higher the depth of a node, the higher the entropy of its embedding. Interestingly, the low-rank Gaussian embeddings achieve the highest correlations, indicating that they are better in preserving depth information.

Last, we compared the performance of the diagonal and rank-1 Gaussian embeddings when using the same number of parameters per node. The results showed that the performance of both embeddings was similar for preserving the shortest path lengths (Table 6), even though the rank-1 ones were lower-dimensional. Nevertheless, the higher-dimensional diagonal embeddings were unable to preserve node depth information as accurately as the lower-dimensional rank-1 embeddings. This comparative analysis demonstrates again the representational advantages of the low-rank approximation.

**Table 2:**
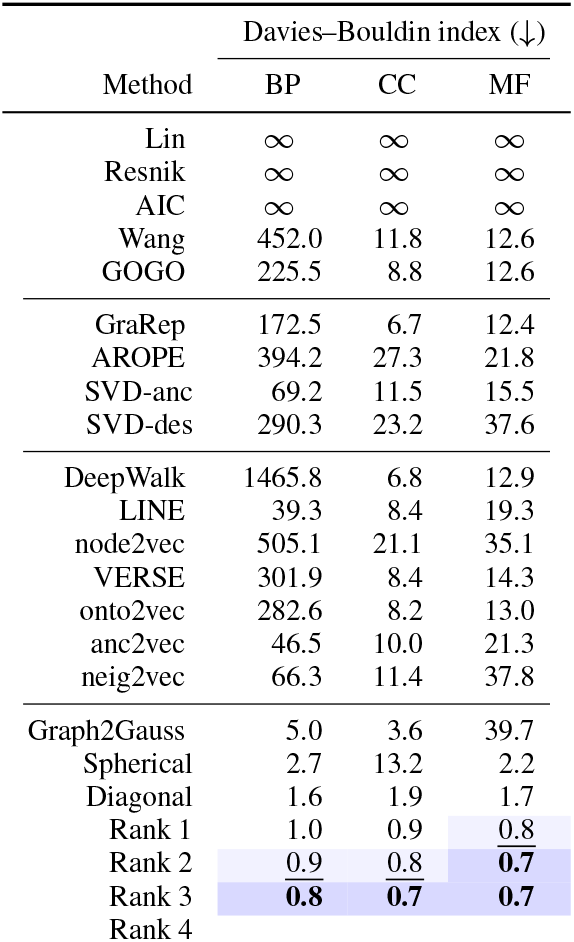
Davies-Bouldin indexes between groups A and D.

**Table 3:**
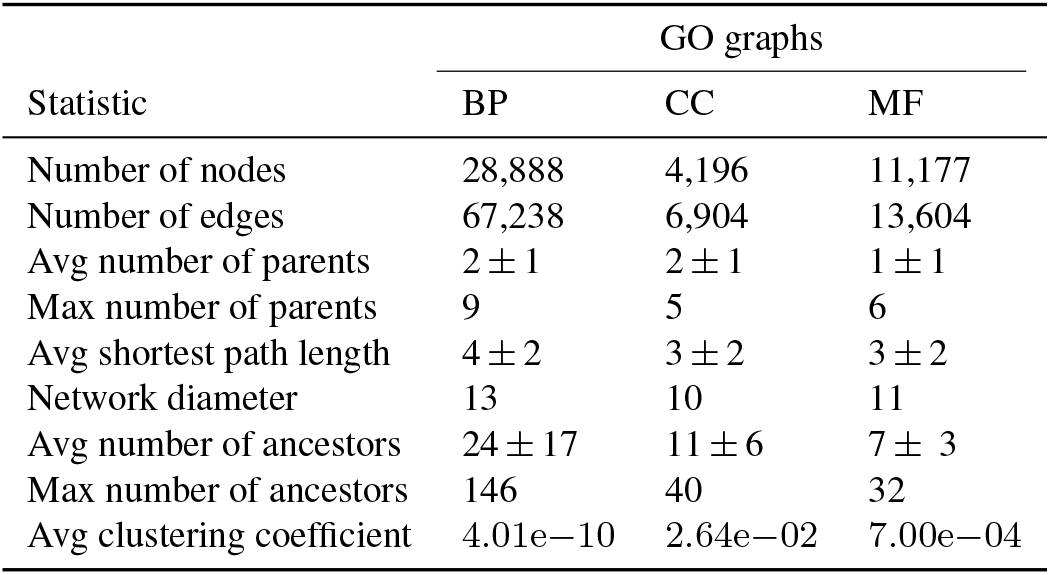
Relevant features of the directed acyclic graphs that compose the GO.

| Statistic | GO graphs |  |  |
| --- | --- | --- | --- |
|  | BP | CC | MF |
| Number of nodes | 28,888 | 4,196 | 11,177 |
| Number of edges | 67,238 | 6,904 | 13,604 |
| Avg number of parents | $2 \pm 1$ | $2 \pm 1$ | $1 \pm 1$ |
| Max number of parents | 9 | 5 | 6 |
| Avg shortest path length | $4 \pm 2$ | $3 \pm 2$ | $3 \pm 2$ |
| Network diameter | 13 | 10 | 11 |
| Avg number of ancestors | $24 \pm 17$ | $11 \pm 6$ | $7 \pm 3$ |
| Max number of ancestors | 146 | 40 | 32 |
| Avg clustering coefficient | $4.01\text{e}-10$ | $2.64\text{e}-02$ | $7.00\text{e}-04$ |

**Table 4:** Time differences between approximations of covariance matrices.

| Model comparison |  | Fold difference |  |  |
| --- | --- | --- | --- | --- |
|  |  | BP | CC | MF |
| Diagonal | Spherical | 0.99 | 0.99 | 0.99 |
| Rank 1 | Diagonal | 3.75 | 2.98 | 3.74 |
| Rank 2 | Rank 1 | 1.61 | 1.45 | 1.59 |
| Rank 3 | Rank 2 | 1.13 | 1.12 | 1.14 |
| Rank 4 | Rank 3 | 1.09 | 1.08 | 1.09 |

**Table 5:** Correlation between the alternative definition of node depth and embedding entropy.

| Model | Pearson correlation (↑) | | |
| --- | --- | --- | --- |
|  | BP | CC | MF |
| Spherical | $0.16 \pm 0.08$ | $-0.18 \pm 0.11$ | $-0.04 \pm 0.17$ |
| Diagonal | $0.26 \pm 0.04$ | $0.47 \pm 0.03$ | $0.46 \pm 0.03$ |
| Rank 1 | **$0.34 \pm 0.01$** | **$0.58 \pm 0.03$** | $0.54 \pm 0.02$ |
| Rank 2 | $0.30 \pm 0.02$ | $0.57 \pm 0.03$ | **$0.62 \pm 0.01$** |
| Rank 3 | $0.27 \pm 0.01$ | $0.52 \pm 0.01$ | $0.61 \pm 0.01$ |
| Rank 4 | $0.30 \pm 0.03$ | $0.54 \pm 0.01$ | $0.59 \pm 0.03$ |
| Graph2Gauss | $-0.13 \pm 0.08$ | $-0.13 \pm 0.11$ | $-0.02 \pm 0.01$ |

**Table 6:** Performance of representations with the same number of parameters per node. Significant correlations are boldfaced.

| Feature | Graph | Pearson correlation (↑) | | | | | |
| --- | --- | --- | --- | --- | --- | --- | --- |
|  |  | 30 params per node |  | 40 params per node |  | 50 params per node |  |
| | | Diag ($d=30$) | Rank1 ($d=15$) | Diag ($d=40$) | Rank1 ($d=20$) | Diag ($d=50$) | Rank1 ($d=25$) |
| Path length $>$ | BP | **$0.90 \pm 0.01$** | $0.88 \pm 0.01$ | $0.91 \pm 0.01$ | $0.90 \pm 0.01$ | $0.91 \pm 0.01$ | $0.90 \pm 0.01$ |
| | CC | **$0.85 \pm 0.01$** | $0.79 \pm 0.02$ | **$0.87 \pm 0.01$** | $0.81 \pm 0.02$ | **$0.86 \pm 0.02$** | $0.81 \pm 0.03$ |
| | MF | $0.88 \pm 0.01$ | $0.87 \pm 0.01$ | $0.88 \pm 0.01$ | $0.87 \pm 0.01$ | **$0.89 \pm 0.01$** | $0.86 \pm 0.01$ |
| Path length $<$ | BP | $0.93 \pm 0.01$ | $0.93 \pm 0.01$ | $0.93 \pm 0.01$ | $0.93 \pm 0.01$ | $0.93 \pm 0.01$ | $0.94 \pm 0.01$ |
| | CC | $0.90 \pm 0.01$ | $0.90 \pm 0.02$ | $0.89 \pm 0.01$ | $0.91 \pm 0.02$ | $0.91 \pm 0.02$ | $0.91 \pm 0.02$ |
| | MF | $0.93 \pm 0.01$ | **$0.96 \pm 0.01$** | $0.93 \pm 0.01$ | **$0.96 \pm 0.01$** | $0.93 \pm 0.01$ | **$0.96 \pm 0.01$** |
| Node depth | BP | $0.66 \pm 0.02$ | **$0.73 \pm 0.03$** | $0.63 \pm 0.02$ | **$0.71 \pm 0.01$** | $0.59 \pm 0.02$ | **$0.70 \pm 0.02$** |
| | CC | $0.63 \pm 0.02$ | **$0.70 \pm 0.02$** | $0.64 \pm 0.01$ | **$0.69 \pm 0.01$** | $0.62 \pm 0.02$ | **$0.69 \pm 0.03$** |
| | MF | $0.62 \pm 0.05$ | **$0.67 \pm 0.05$** | $0.58 \pm 0.02$ | **$0.68 \pm 0.01$** | $0.63 \pm 0.02$ | **$0.68 \pm 0.02$** |

### 4.4 Ablation study on the loss components

We conducted an ablation study to gain a deeper understanding on the contribution of the three components of the loss function shown in Eq.2. To this aim we used gGN to learn node embeddings, from each real-world graph, by turning off one of the three loss components during the whole training process. The resulting Gaussian embeddings were then used to calculate how well correlated they were with respect to the structural features analyzed in §4.3. Next, the correlations obtained from the ablated embeddings were compared with those obtained from the original ones, which were previously reported in Table 1. We expected that if a loss component had a role in capturing one of the structural features, the embeddings learned when this component is turned-off should show a lower correlation in comparison to the original one. The three covariance approximations studied here (spherical, diagonal and low-rank) were included in this approximation analysis.

Figure 4 visually compares the Pearson correlations obtained from the original embeddings (x-axis) and the ablated ones (y-axis). Rows show the evaluated three graph features whereas each column indicates which loss component was ablated. Therefore, each dot indicates the correlation values obtained by one of the Gaussian embeddings (spherical, diagonal and low-rank) learned from a particular graph when a given loss component is ablated or not. Since the main diagonal represents cases where ablated and original embeddings obtain the same correlations, dots falling below the main diagonal indicate that the correlations obtained by the original embeddings were better (higher) than the ablated ones. The results show that when the loss component 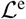 is ablated, almost all the embeddings face difficulties in correctly capturing the three graph features, and particularly in preserving node depths. This is expected as this loss component is preserving the local structures around each node. Next, when ablating the loss components 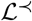 and 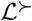, moderate negative impacts are observed on capturing the shortest path length with the normal (≺) and the inverted (≻) directions, respectively. This demonstrates the role of these two components in preserving the shortest path lengths between nodes. Moreover, note that ablating a loss component may have impact on a specific structural feature but not on others, reflecting the relevance of each loss component in capturing specific graph information. For example, ablating the loss component aimed at capturing the shortest path lengths of descendants 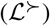 has almost no impact on capturing shortest path lengths of ancestors (≻) but a mild impact on capturing node depths. All these results indicate that the three loss components are needed for learning Gaussian embeddings capable of properly preserving meaningful graph features.

**Figure 4:**
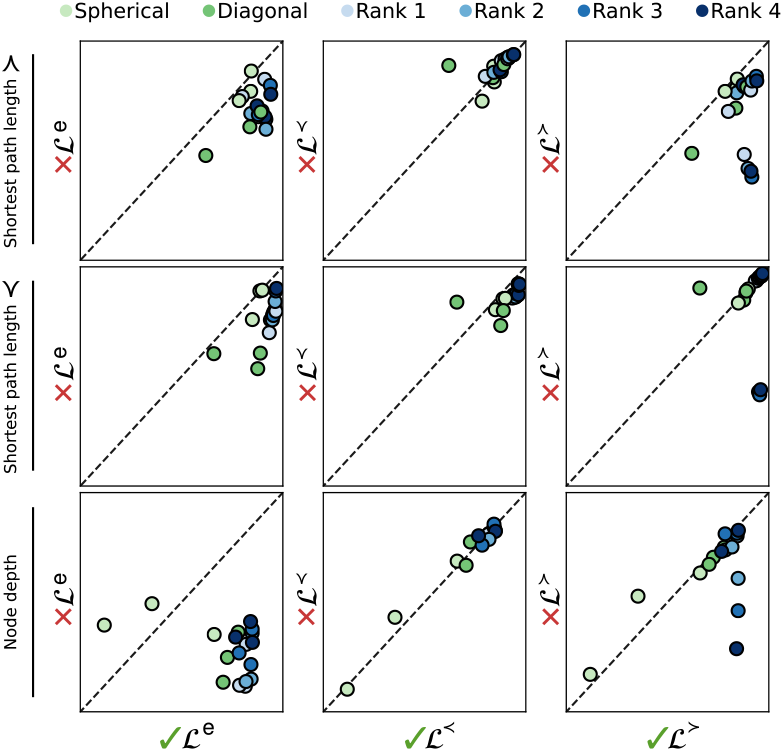
Ablation study to assess each individual loss component.

### 4.5 KL divergence for measuring semantic similarity

Since the KL divergence between the distributions encoded by the embeddings captures the length of the shortest path between the corresponding nodes, we further analyzed whether this divergence could be used as a way of measuring the semantic similarity between nodes [44, 45, 46]. This task is fundamental for comparing the functional relatedness of gene products based on their annotated GO terms, and lies at the heart of automating annotation of protein functions [47, 48, 49].

Given a graph, we evaluated whether the KL divergences (i.e., semantic similarities) calculated from a particular node *i* to all the others were able, on their own, to discriminate between the ancestors (A) and descendants (D) of *i*. By performing this systematically for all nodes, we ultimately measured whether the resulting semantic similarities could discriminate between the groups A and D. To quantify such discrimination, the Davies-Bouldin index [50] was used, as it measures the level of overlapping between two distributions. The lower the Davies-Bouldin index is, the less overlapped two distributions are. Thus, lower indices indicate better discriminations between the groups A and D. In this evaluation, we included the well-known methods used for measuring semantic similarities, as well as methods that learn node representations, which calculate semantic similarities through cosine similarities between embeddings. More experimental details are provided in §E.

Table 2 shows the Davies-Bouldin indexes calculated from the semantic similarities obtained for each method on each real-world graph. In general, methods not using distributions as node representations show important difficulties (high indexes) to discriminate between ancestors and descendants. For example, the indexes of Lin, Resnik and AIC are infinite on the three graphs, indicating a complete overlap between the semantic similarities of ancestors (group A) and descendants (group D). This is expected since they calculate the semantic similarity between two nodes based on their common ancestors [23, 24, 51]. Better discriminations between both groups (low indices) are obtained by methods that learn node representations, such as LINE and onto2vec, but in particular those that learn Gaussian distributions. In the latter, the performance of gGN is the best one, even when using diagonal covariance matrices. Notably, the best indices are obtained by the low-rank approximations, and they get better as the rank is higher. Taken together, these results clearly demonstrate not only the advantages of using the proposed Gaussian representations but also that the KL divergence is a powerful approach for measuring the semantic similarity between nodes.

## 5 Conclusion

This work presents gGN as a novel representation for graph nodes based on low-rank Gaussian distributions. The main benefits of the proposed approach stems from not approximating covariance matrices with diagonal matrices as is usually done. Instead, our proposal uses low-rank covariance matrices that, unlike diagonal ones, are capable of representing correlations between the dimensions of an embedding space. We demonstrated here that such correlations are crucial for expressing hierarchical structures in knowledge graphs. Since estimating non-diagonal covariance matrices has a quadratic cost, we used a low-rank approximation to make such estimation computationally affordable. In addition, a novel loss function is introduced to learn in an unsupervised manner the low-rank representations such that they preserve both node depths and asymmetric relationships between nodes. Empirically, our results show that the proposed representation is better than existing approaches in capturing hierarchical structures and in semantics similarity tasks.

A promising direction for future work is learning low-rank node representations using an alternative input matrix for the lengths of the shortest paths between nodes. Classically, such matrices are calculated with Dijkstra’s and Floyd-Warshall’s algorithms. Even though the matrix of lengths is calculated only once for each graph in our approach, the aforementioned algorithms are limited to relatively small networks due to their high computational complexity, which is *O*(*n*^3^) time. Nevertheless, both algorithms are parallelizable and there exist GPU implementations [52, 53]. Alternatively, approximate methods with acceptable accuracy have been also proposed for scaling up computations to graphs with millions of edges [54, 55]. Among them, a promising approach is the so-called landmark-based methods [54]. It selects a fixed set of landmark nodes from which their shortest paths to all other nodes are precomputed. By exploiting geometric properties among landmarks, one can approximately compute the shortest path between any two nodes, with a cost in the order of the number of landmarks.

## Appendix

### A KL divergence between low-rank Gaussian distributions

Let 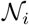 and 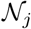 be two multivariate Gaussian distributions 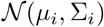 and 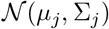 where 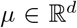 and 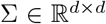 are the mean and covariance matrix, respectively. Let assume that each covariance matrix is represented by a low-rank form

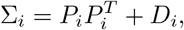

where *D_i_* is a diagonal matrix in 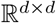 and 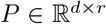. The (reverse) KL divergence between 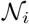 and 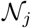 has the following closed form [37]

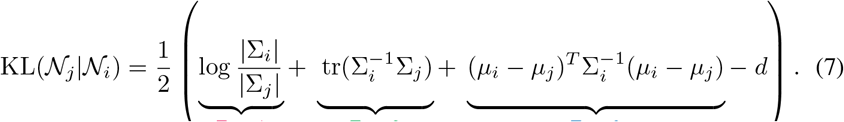

Its computational cost can be substantially reduced by exploiting the low-rank forms of the first three terms.

#### A.1 Calculating KL Term 1

This term basically involves calculating the determinant of a covariance matrix Σ. The determinant of a low-rank form is

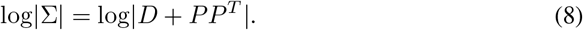

This form enables us to use the matrix determinant lemma

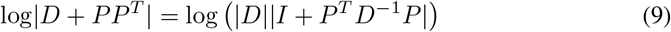

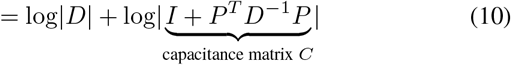

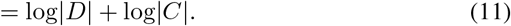

Here, 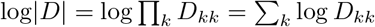. On the other hand, by using the Cholesky decomposition, the determinant of the capacitance matrix 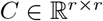 is

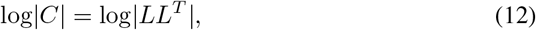

where 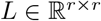 is a lower triangular matrix. Here, calculating the Cholesky decomposition takes *O*(*r*^3^) time, where *r* ≪ *d*. Thus, the determinant can be calculated as follows

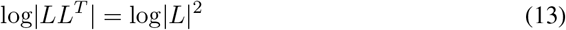

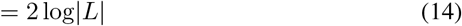

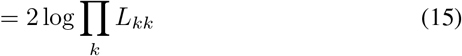

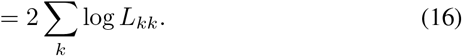

Taken together, **Term 1** can be expressed as

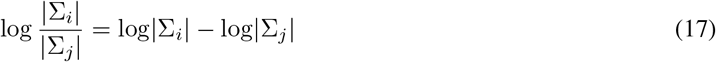

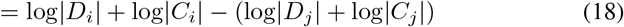

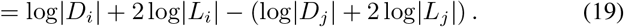

This formula shows that computing the determinant of both covariance matrices boil down to computing the determinants of diagonal and triangular matrices, as simply the product of their diagonal values.

#### A.2 Calculating KL Term 2

Now, let us see how to compute the second term:

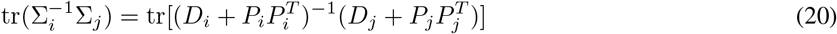

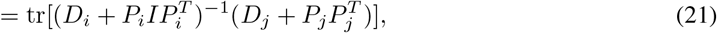

using the Woodbury matrix identity, the first factor can be re-written

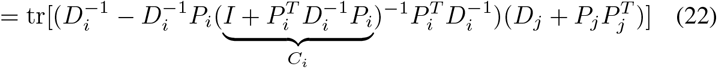

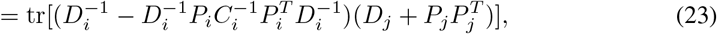

now, we can re-used the Cholesky decomposition of *C_i_*

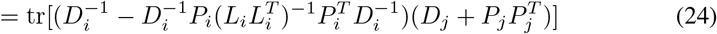

by algebraically manipulating the factorization of *C_i_*, a symmetric structure (*A*) can be exposed

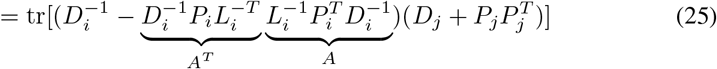

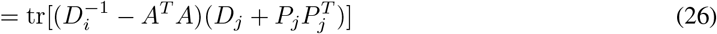

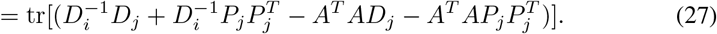

Due to the linearity of the trace operator, the latter equation is equivalent to

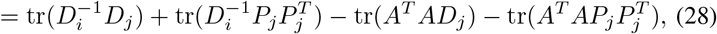

to further simplify this expression, we can use the Cholesky decomposition of some diagonal matrices

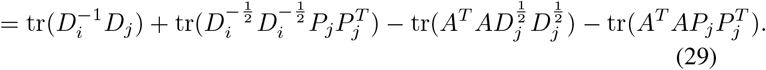

By using the cyclic property of the trace operator

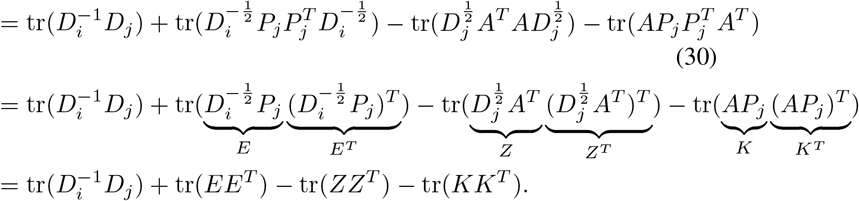

Therefore, this KL term is reduced in applying the trace operator on four matrix multiplications. However, since the matrix multiplications involve transposed matrices, operations can be further reduced. Note that 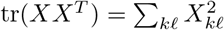.

#### A.3 Calculating KL Term 3

Since the difference between means can be thought of as a displacement Δ, we can use it to re-write the third term for making the notation easier

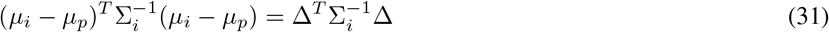

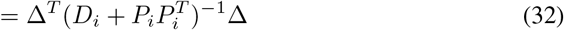

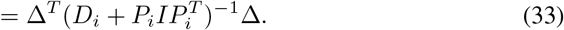

The intermediate factor can be re-expressed using the Woodbury matrix identity

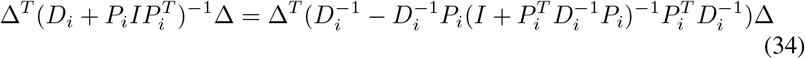

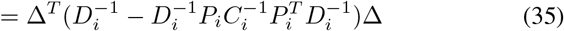

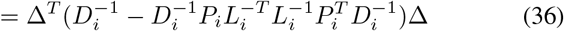

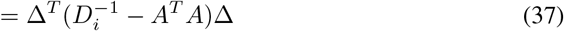

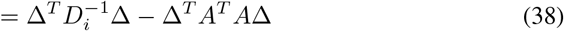

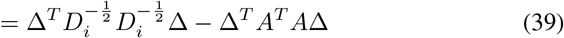

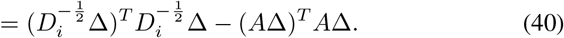

This last line is cheap to compute as it involves diagonal matrices and the matrix *A*, which was previously computed for KL Term 2.

### B Data and training details

For the experiments, we used the three directed acyclic graphs obtained from the GO [7] (release 2020-10-06). Each was constructed from the *is-a* relationships defined by one of the three sub-ontologies: BP (Biological Process), CC (Cellular Component) and MF (Molecular Function). Relevant features of these graphs are shown in Table 3. For each graph, a matrix *S* containing the shortest path lengths was constructed using the Floyd-Warshall algorithm as provided by SciPy [56]. The resulting matrix S was then used as input to gGN (https://github.com/blindcosmos/ggn) to learn the node embeddings by using three different seeds, for analyzing learning variability.

Although all technical details about gGN are available in its code, it is worth mentioning the following points. To learn each embedding 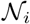, the mean *μ_i_* and covariance factor *P_i_* were initialized randomly using a normal distribution with mean 0 and variance 1, while the diagonal matrix *D_i_* was initialized with 1s, following previous recommendations [14]. Since each covariance matrix Σ_*i*_ needs to be positive definite (|Σ_*i*_| > 0), the diagonal matrix *D_i_* was clipped to lie within the hypercube [0.01, ∞]^*d*^ each time before computing the loss function. This lower upper bound was chosen through numerous experimental analyses that demonstrated that such value made the learning process very stable by preventing the determinants of the covariance matrices from reaching zero. In all cases, the training process was performed during 1,000 epochs using batches of 128 random nodes and the Adam optimizer [57] with default parameters. Following standard methodologies [32, 58], no test and validation sets were used since the aim here is to learn node embeddings that completely fit the graph structure. All experiments were run on an i7-5960X processor equipped with eight 3-GHz dual cores in a server with 64GB of RAM and a single Nvidia Titan X GPU.

In all the experiments involving comparison among the different Gaussian distributions (i.e., spherical, diagonal and low-rank), all the embeddings were learned by using latent spaces with the same dimensionality. This is because all models are single-layered without activations and thus, an attempt to set an equal number of parameters per node would directly change the space dimensionality. For instance, a less expressive Gaussian model (e.g., diagonal) with a latent space with more dimensions than the space of a more expressive Gaussian model (low-rank) can prevent us from determining whether differences on model performance are due to space dimensionality or Gaussian types.

### C Time complexity of learning Gaussian embeddings

The time curves in Figure 5 show the time spent by gGN when learning embeddings using different covariance matrices on the three graphs evaluated here. When comparing the curves among them, it can be seen that the higher time consumption occurs on BP followed by MF, with CC displaying the lowest times. This is expected as the time complexity linearly depends on the number of nodes, and BP and MF have the largest number of nodes (Table 3). Regardless of the graph, the curves also show that the lowest computational cost is obtained by the spherical and diagonal approximations, with essentially the same time complexity since they have the same number of parameters.

**Figure 5:**
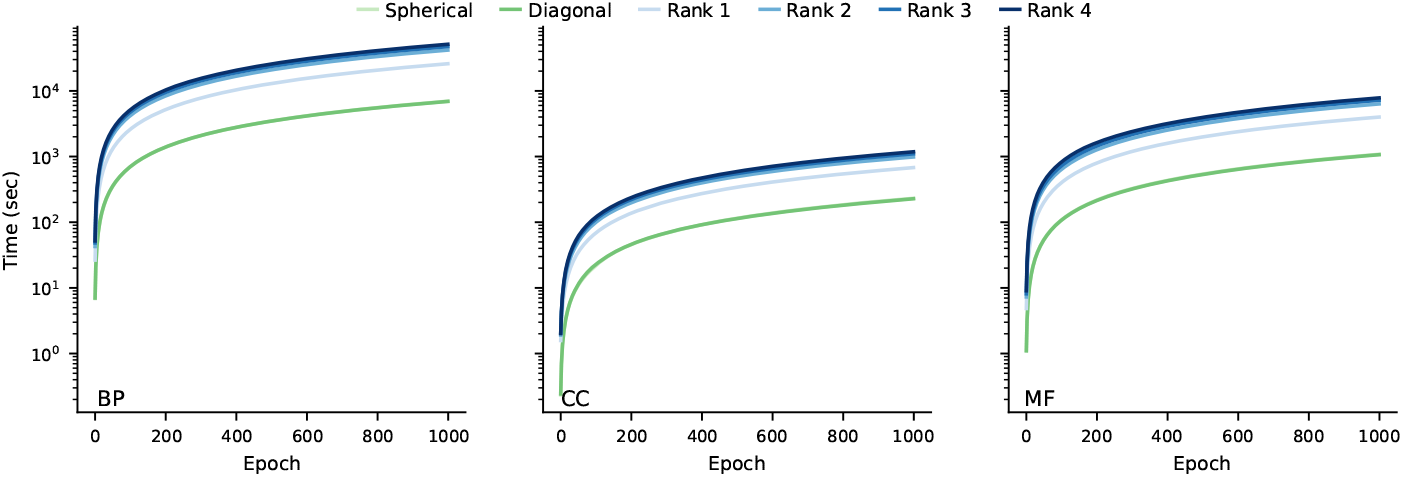
Time complexity curves. Time spent by gGN when learning 10-dimensional Gaussian embeddings with different types of covariance matrices.

The next lowest times are obtained by the rank-1 approximations. The higher the rank is, the higher the time complexity is, which is expected as the time complexity grows as a function of the rank.

Table 4 quantitatively shows how many folds higher it is the time consumed by a model with respect to another. We can see more clearly that the time complexity of spherical and diagonal embeddings are almost the same. Among the low-rank approximations, the time spent by the rank 2 is 1.6-fold higher than the rank 1, and this time difference between ranks tends to reduce as both consecutive ranks are higher.

We also investigated how the time complexity scales as a function of the graph size by comparing the time spent to learn the Gaussian models on 1000 epochs. The results show that the time increases as the graph size increases (Fig. 6). This result also showed that, regardless of the type of Gaussian model, the time complexity per-node is proportional to the number of nodes of the graph.

**Figure 6:**
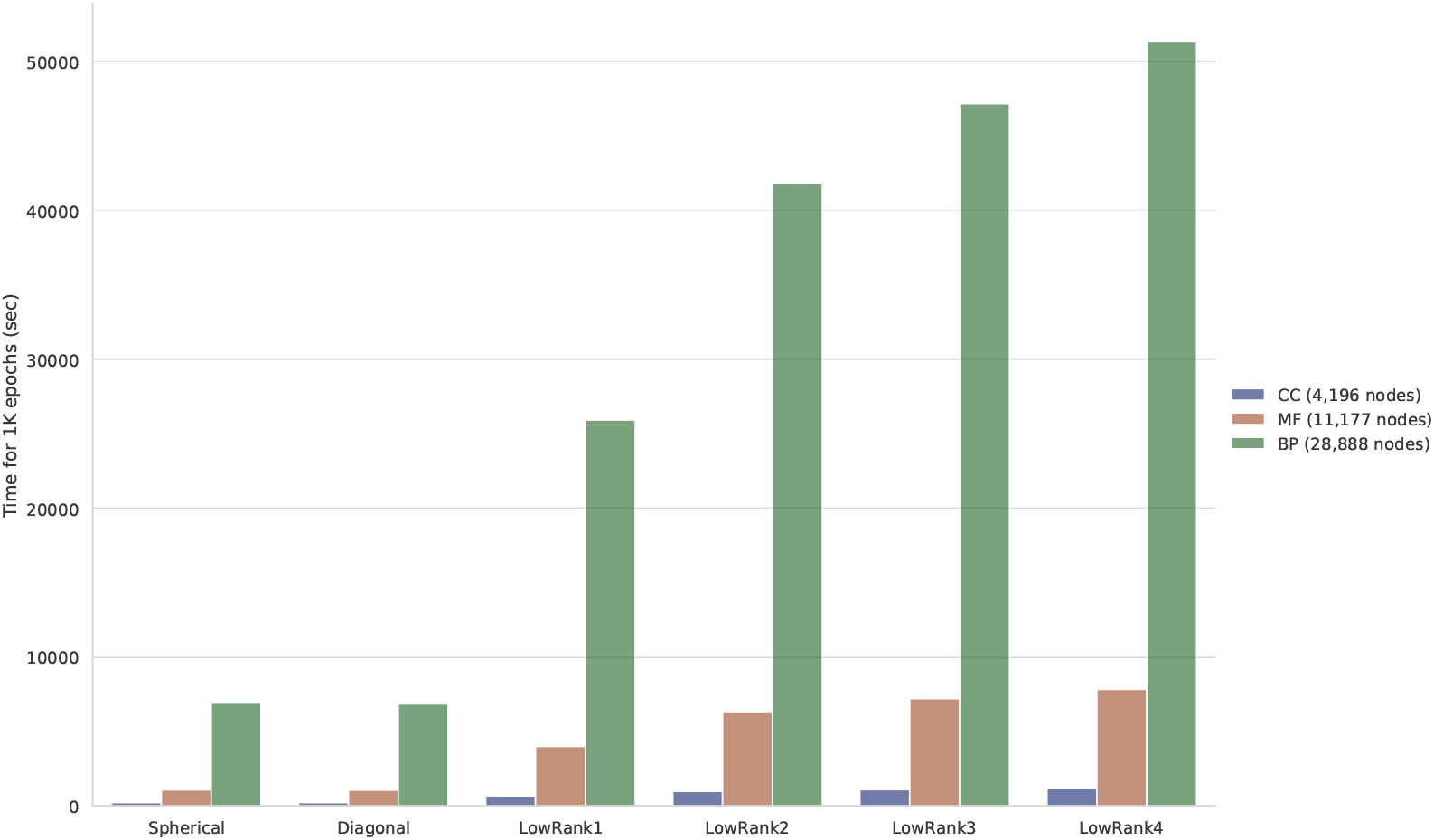
Time complexities in function of the graph.

### D Graph features preserved by Gaussian embeddings

We evaluated whether the Gaussian embeddings were preserving both the shortest path lengths between nodes and the depth of nodes. To this aim, we computed the KL divergence between Gaussian embeddings and then we compared it against the shortest path lengths between the corresponding nodes (§D.1). Similarly, to evaluate that the node depths were preserved, we calculated the entropy of each Gaussian embedding and then we compared it against the depth of its node (§D.2). Since the KL divergence and entropy are not normalized, hampering comparative analyses through visualizations, both measures were normalized for each type of embedding (spherical, diagonal and low-rank) on each graph. More concretely, given the KL divergences (or entropies) calculated from embeddings learned from a graph, these values were normalized by subtracting the minimum value and then dividing them by the difference between the maximum and minimum values.

#### D.1 KL preserves shortest path length

Figure 7 shows the resulting KL divergences between embeddings when grouped according to the shortest path lengths between the nodes, which range from 1 to 13 (BP), 1 to 10 (CC) and 1 to 11 (MF). Each group is showing the distribution of KL divergences calculated between embeddings representing pairs of nodes sharing the same shortest path lengths. Such distributions are depicted as violins and their different quartile values as boxes. The results show that for all embeddings the KL divergence is positively correlated with the shortest path length. This was quantified more precisely by calculating how linearly correlated both variables were and plotted as red dashed lines. The highest correlations were notably achieved by the low-rank embeddings, indicating that they are properly preserving the graph topology.

**Figure 7:**
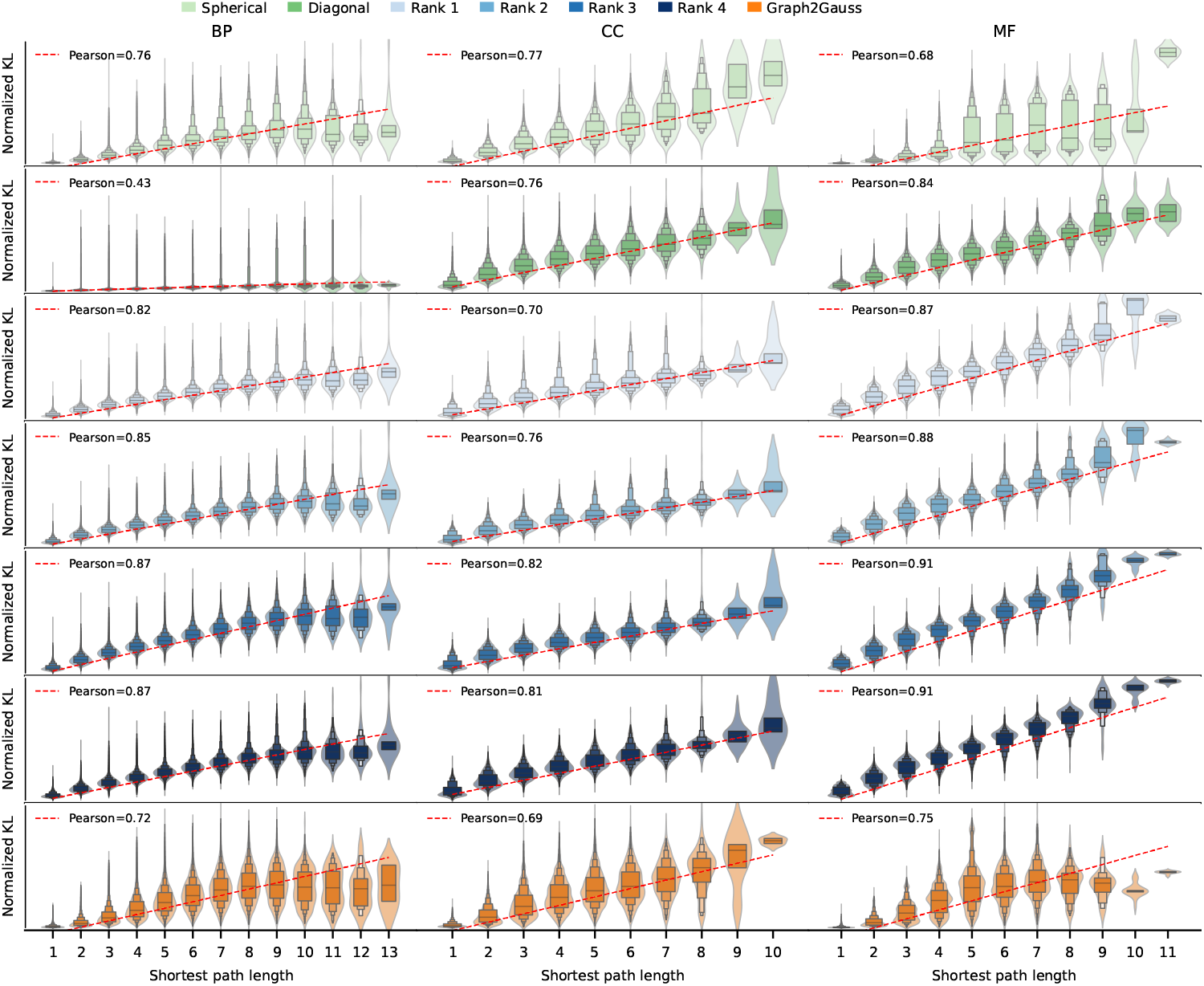
Shortest paths lengths (≻) between nodes and KL divergence between their Gaussian embeddings.

Since the KL divergence is asymmetric, we also evaluated whether the previous positive correlation was also held when inverting the directions of all edges in each graph. The results are shown in Figure 8. As expected, it shows that the KL is strongly correlated with the length of the shortest paths obtained from inverting directions. In particular, the highest correlations are obtained by the low-rank embeddings.

**Figure 8:**
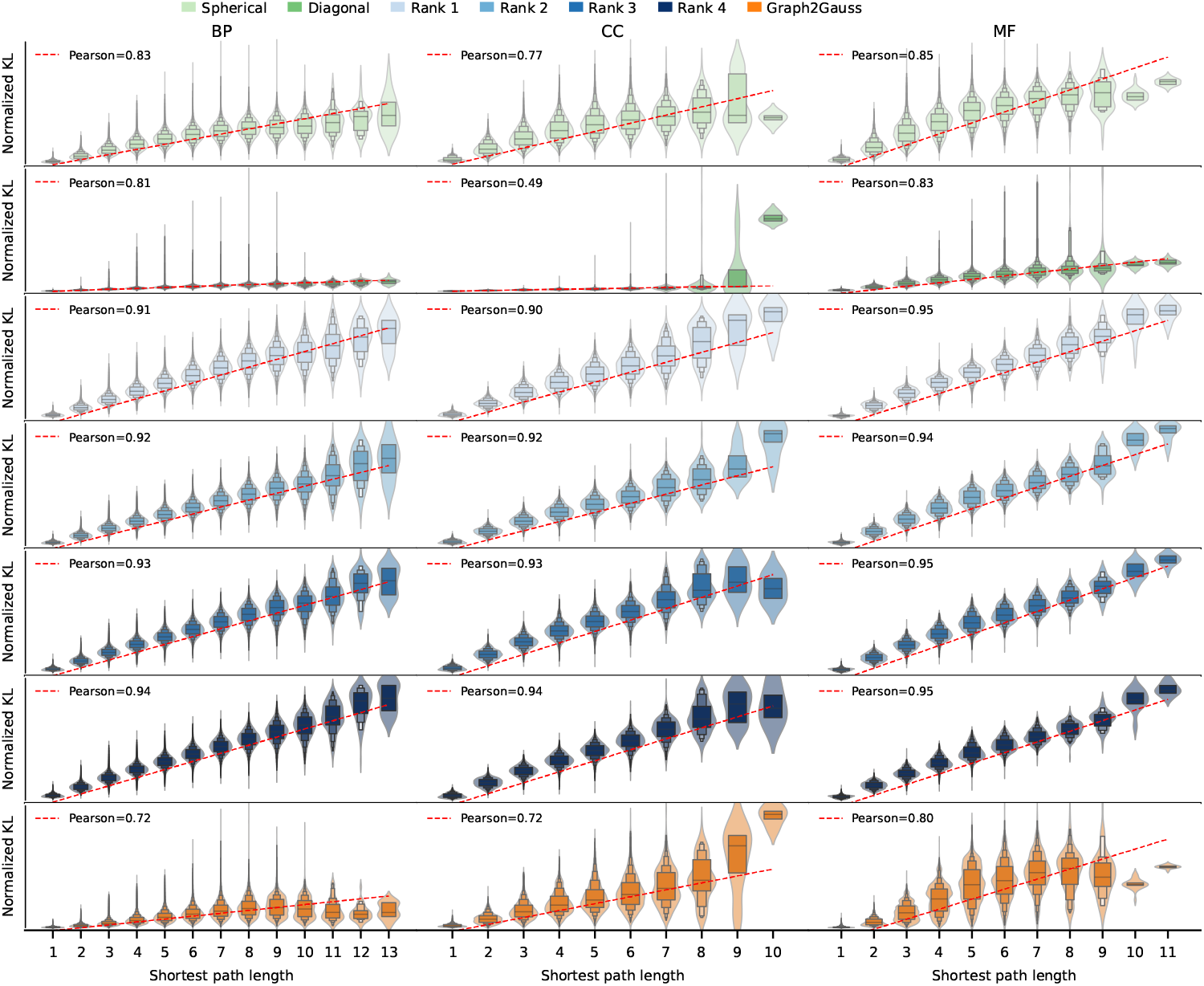
Shortest paths lengths (≺) between nodes and KL divergence between their Gaussian embeddings.

#### D.2 Entropy preserves node depth

To this evaluation, we defined the depth of a node as its number of ancestors. To calculate the entropy, we took advantages that embeddings are Gaussian distributions and used the following closed form formula:

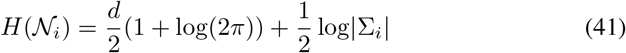

where |·| is the determinant. Figure 9 shows the results of comparing the node depths with the entropies of their corresponding embeddings. Entropies calculated from nodes sharing the same depth are grouped and represented as violins, whereas the quantiles are depicted as boxes inside the violins. In almost all embeddings, the node depth is positively correlated with the entropy. However, this correlation is not observed for the spherical embeddings on CC and MF. This is expected as the spherical embeddings can only represent overly limited density shapes, which seem to be not sufficient for properly encoding node depths through patterns of encapsulation between embeddings. The level of correlation was quantitatively measured by calculating how linearly correlated the depth and the entropy were for each method on each graph. The resulting correlations are shown as red dashed lines. The highest correlations are achieved by the low-rank embeddings, indicating that they are better suited for preserving depth information.

**Figure 9:**
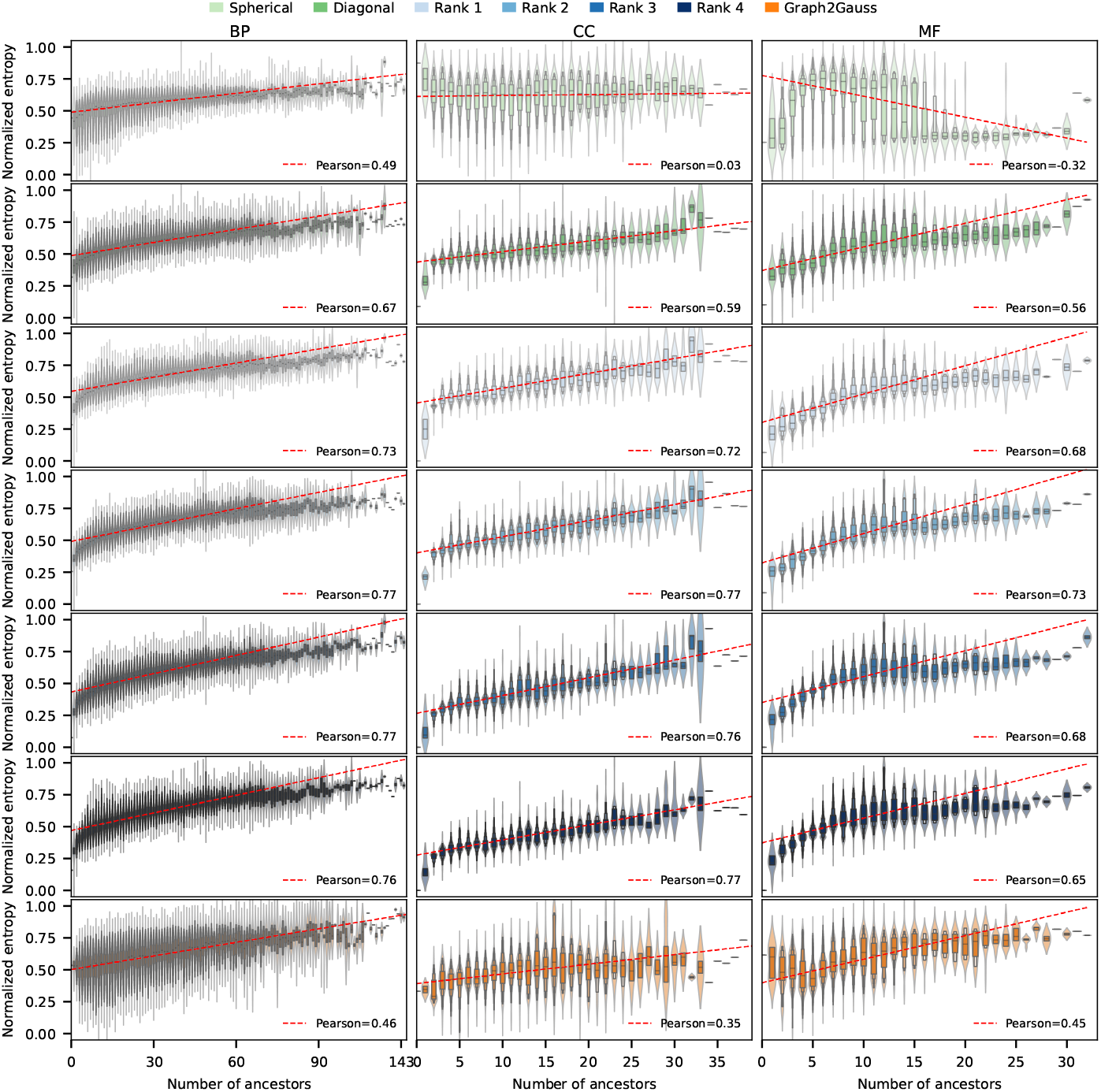
The number of ancestors of a node and the entropy of its corresponding Gaussian embedding.

Since the graphs used in the experiments are DAGs, we defined the node depth alternatively as the length of the shortest path between a node and its root. Next, we measured how correlated this new definition was with the embedding entropy. The results are shown in Figure 10. Here, we can see that the low-rank embeddings achieve the highest correlations, as indicated by the linear correlations plotted as red dashed lines. Table 5 shows these results more clearly by listing all the Pearson correlation coefficients obtained for each method on each graph. This high correlation achieved by the low-rank embeddings demonstrates that depth information is successfully preserved.

**Figure 10:**
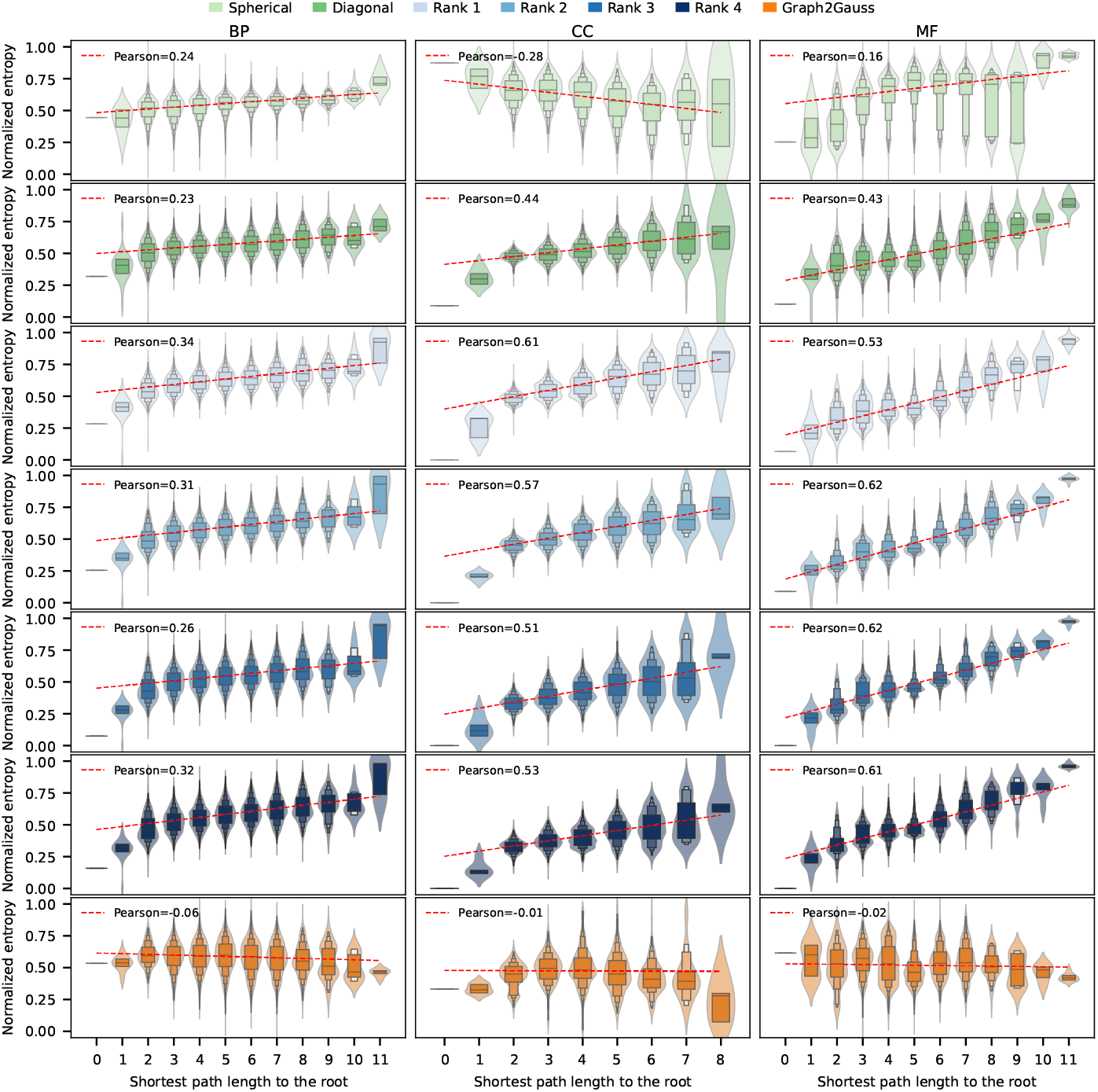
The shortest path length of a node to the root and the entropy of its corresponding Gaussian embedding.

In addition, we compared the diagonal and rank-1 embeddings when using the same number of parameters per node. For representing a graph node, a diagonal embedding of dimension *d* uses the same number of parameters than a rank-1 embedding of dimension d/2. This comparison evaluated the performance of both embeddings on preserving shortest path lengths and node depths. The results are shown in Table 6. In general, the performance of the diagonal and rank-1 embeddings is similar. Nevertheless, the low-rank embeddings are better than the diagonal ones on preserving node depths, even when the latter used the double of dimensions.

### E Semantic similarity task

To evaluate node representations on the task of measuring semantic similarity between nodes, we extracted all possible pairs of nodes of a given directed graph and then labeled them according to the relationships held between both nodes in each pair. More concretely, for a given node *i*, the other nodes in a directed graph can be partitioned into three groups: ancestors, descendants or neither of them, as is schematically shown in Figure 11. Hence, each pair between node i and node *j* can be labeled into one of these three groups. Therefore, the task consisted on analyzing whether the semantic similarities calculated from the (Gaussian) representations of two nodes were able, on their own, to discriminate among the three groups of node pairs.

**Figure 11:**
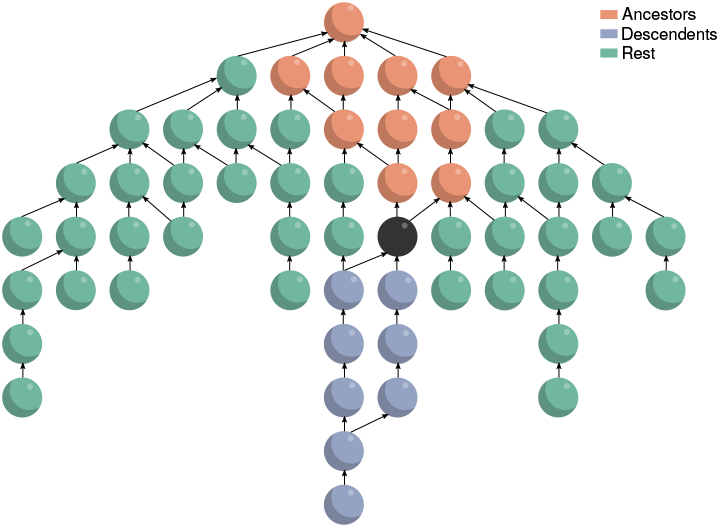
Groups of node pairs based on graph partitioning. Given any node (black), it can be used as a reference to partition a directed graph into three disjoint sets of nodes: its ancestors, its descendants and the rest of the nodes.

#### E.1 Selected methods for the semantic similarity task

For the task of measuring semantic similarity between nodes, we selected a number of representative baseline methods. Some of them were specifically designed for this task while others are well-known methods that have been proposed for learning node representations from graphs. Table 7 lists the selected methods that are grouped into five wide categories according to their computational approach, indicated in the column Category. The next columns indicate corresponding year and repository. On top, the classic methods are Lin [23], Resnik [24], AIC [51], Wang [59], and GOGO [60]. They stem from ideas based on information theory previously applied on the WordNet hierarchy. The other four categories exclusively include methods that learn node representations from a given graph.

**Table 7:** Methods used on the task of measuring semantic similarity.

| Method | Category | Year | Repository |
| --- | --- | --- | --- |
| Lin | Classic | 1998 | - |
| Resnik | Classic | 1999 | - |
| AIC | Classic | 2013 | - |
| Wang | Classic | 2007 | https://github.com/tanghaibao/goatools |
| GOGO | Classic | 2018 | https://github.com/zwang-bioinformatics/GOGO |
| GraRep | Matrix factorization | 2015 | https://github.com/benedekrozemberczki/GraRep |
| AROPE | Matrix factorization | 2018 | https://github.com/ZW-ZHANG/AROPE |
| SVD-anc | Matrix factorization | - | - |
| SVD-des | Matrix factorization | - | - |
| DeepWalk | Random walk | 2014 | https://github.com/phanein/deepwalk |
| LINE | Random walk | 2015 | https://github.com/tangjianpku/LINE |
| node2vec | Random walk | 2016 | https://github.com/aditya-grover/node2vec |
| VERSE | Random walk | 2018 | https://github.com/xgfs/verse.git |
| onto2vec | Neural network | 2018 | https://github.com/bio-ontology-research-group/onto2vec |
| anc2vec | Neural network | 2022 | https://github.com/sinc-lab/anc2vec |
| neigh2vec | Neural network | 2022 | https://github.com/sinc-lab/anc2vec |
| Graph2Gauss | Gaussian | 2017 | https://github.com/abojchevski/graph2gauss |

In the matrix factorization category, the methods selected are GraRep [27], AROPE [61], and DeepWalk [28]. They first define a high-dimensional matrix in which structural features of each node are described. Next, the dimensionality of this matrix is reduced in order to obtain low-dimensional (dense) representations of nodes. This group includes two additional methods named SVD-anc and SVD-des that are specially prepared for the semantic similarity task (technical details of these methods are provided in §E.2).

The next category contains methods based on random walks: LINE [29], node2vec [30], and VERSE [31]. These methods perform random walks starting from a given node to define its neighborhood stochastically, in order to use it in a self-supervised learning task for building the representation of the node. Here, methods generally follow very tightly the self-supervised objectives proposed by word2vec [32], such as the skip-gram objective where the aim is to predict the neighborhood of a given node by using noise contrastive estimation [62]. LINE optimizes similarities between pairs of node embeddings such that first and second order proximities are preserved. Instead, VERSE builds node embeddings such that the similarities between nodes have low KL divergence with respect to the dot products between their corresponding embeddings [31].

The neural network category includes: onto2vec [58], anc2vec [63], and neigh2vec [63]. These methods that use diverse neural networks architectures aimed at building node representations through self-supervised objectives proposed for capturing relevant structural features of a graph. Finally, the last category includes a method that builds node representations using Gaussian distributions whose covariance matrices are approximated by diagonal matrices instead of low-rank matrices like gGN.

#### E.2 Training details for learning node embeddings

For all classic methods, we used default parameters as proposed by their authors. All methods that learned embeddings used 200 dimensions. For GrapRep, 20 orders were used, each of them with 10 dimensions. For AROPE, default parameters were used and 49 dimensions for each order. For SVD-anc (and SVD-desc), an r-dimensional embedding was obtained for each node by truncating the left singular matrix U at dimension *r*, and multiplying it by the truncated diagonal matrix *D*: *U*_1:*r*_*D*_1:*k*,1:*k*_. The matrices *D* and *U* were obtained by computing the SVD decomposition [40] *M* = *UDV^T^*; here, *M* is the matrix of ancestors (SVD-anc) or descendants (SVD-desc), which were built from BP, CC and MF, respectively. For DeepWalk, 80 random walks per node with a maximum length of 40 were extracted. To construct embeddings, the extracted walks were treated as sentences, and nodes were treated as words, to be used as input to word2vec (https://github.com/tmikolov/word2vec) using the skip-gram objective with parameters: window=5, min-count=0 and iter=200. For LINE, we used 2 orders, 5 negatives, 100 samples and *ρ* = 0.025. In node2vec, *p* =1 and *q* = 1 were used. We used VERSE with *α* = 0.85 and 3 samples. From the axioms extracted by onto2vec, it was given as input to word2vec using the skipgram objective with parameters: window=5, min-count=0 and iter=200. The node embeddings of anc2vec and neigh2vec were downloaded from their public repositories. For Graph2Gauss, we used as input an adjacency matrix built for each graph to learn node embeddings with default parameters.

#### E.3 Semantic similarity computation

Classic methods define the similarity between nodes by combining structural features from the hierarchical graph and the information contents of nodes that are calculated from an annotation corpus [64]. However, if different corpuses of annotations are used, information contents may differ and thus lead to semantic incongruencies, which can negatively impact downstream tasks. To overcome this, the information content was calculated using the intrinsic technique [65]. The intrinsic information content (IC) of a given term/node *i* is

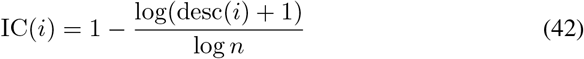

where desc(*i*) is the set of descendants of *i* and *n* is the total number of nodes in the graph.

For the methods that build node representations, we calculated the most commonly used semantic similarity as the cosine [58, 66]. Given two node representations, *v_i_* and *v_j_* in 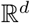, the cosine similarity is

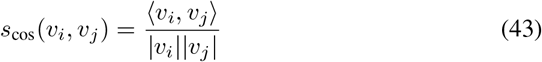

where the dot product of the vectors *v_i_* and *v_j_* is divided by the product of their Euclidean norms. It returns values within the interval [-1, +1], where −1 and +1 indicate total dissimilarity and full similarity, respectively. Since the cosine similarity does not depend on the magnitudes of the vectors, it is totally governed by their angle, and thus *s*_cos_(*i, j*) = *s*_cos_(*j, i*). For Graph2Gauss, the semantic similarity was calculated by using the forward KL divergence for diagonal covariances

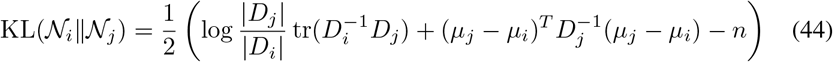

where the determinant log |*D*| = ∑_*k*_ log *D_kk_*, and the inverse of the diagonal matrix is 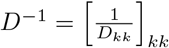.

#### E.4 Results of the semantic similarity task

The semantic similarities obtained for each method are shown in Figure 12 in the y-axis. They are shown by method on the x-axis according to their performance on the ancestors (red), descendants (blue) and rest (green) for the 3 ontologies: BP (top), CC (middle) and MF (bottom). Groups of semantic similarities are plotted using standard boxplots. Because the semantic similarities (KL divergences) calculated from Gaussian embeddings are in an exponential space, due to the loss function used for their optimization, their boxplots are in a log scale to enhance visualization. Similarly, since the obtained semantic similarities vary largely across methods, hampering comparative analyses through visual inspection, they were normalized for each method on each graph. Note that these transformations do not alter the analysis since they do not distort the relative positioning of the semantic similarities obtained by a given method on a particular graph.

**Figure 12:**
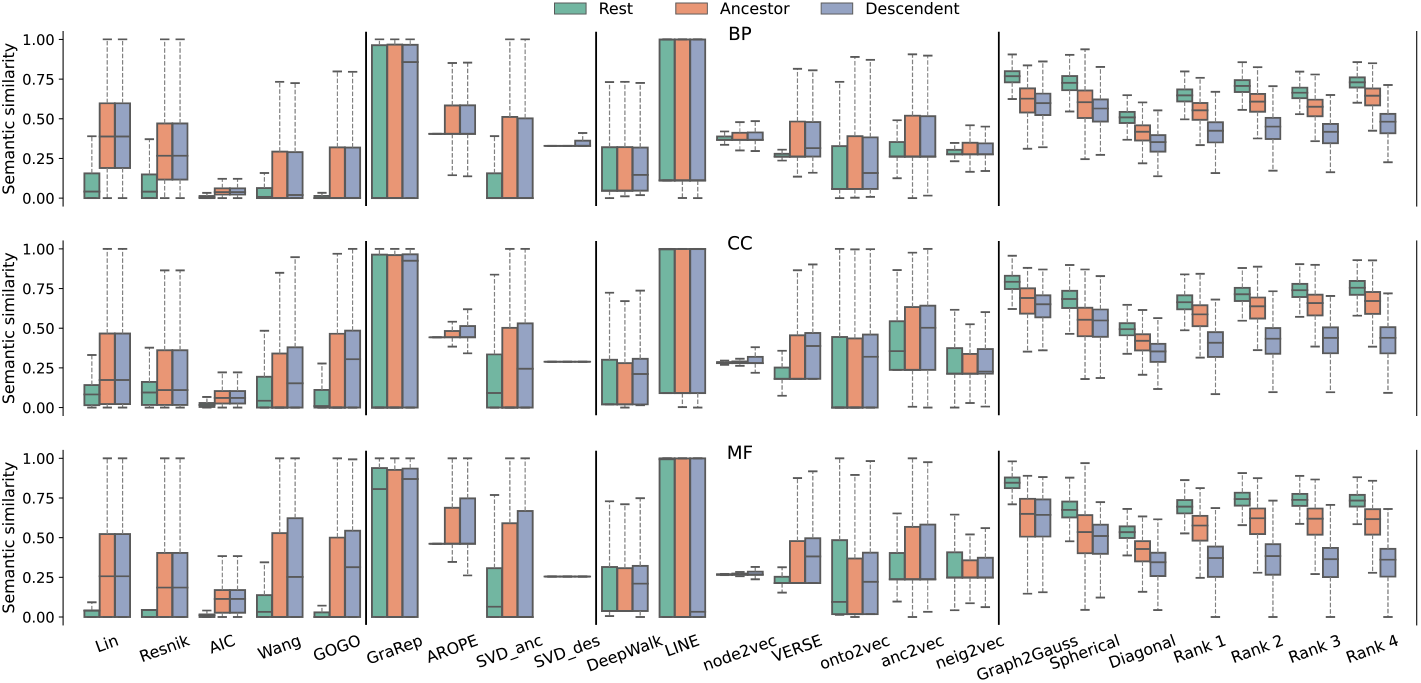
Semantic similarity between pairs of nodes labeled as Ancestors, Descendants and Rest.

If the semantic similarities calculated by a method are good discriminating between two groups (e.g., ancestors and descendants), it is expected for the semantic similarities belonging to these two groups to be poorly overlapped. Since groups of semantic similarities are represented as boxes, no overlapping between boxes is expected, in particular in their most dense regions, which are indicated by the interquartile ranges. Based on this expectation, and analyzing the results in Figure 12, almost all methods show groups overlapped, indicating a poor discriminatory power. Nevertheless, the Gaussian embeddings show higher discriminatory power, since groups are here only slightly overlapped. However, note that this good discriminatory power of the Gaussian embeddings is not general but specific to some embeddings. For example, Graph2Gauss and spherical embeddings show groups of ancestors and descendants with a moderate degree of overlap, whereas the diagonal embeddings show it to a lesser extent. Remarkably, this overlap is not observed among the low-rank embeddings, clearly showing their capability to properly discriminate between groups.

##### E.4.1 Quantifying discriminatory power

To quantify the aforementioned results shown in Figure 12, we used the Davies-Bouldin (DB) index to measure the degree of overlapping between two distributions of semantic similarities. Given two clusters *C_i_* and *C_j_*, the Davies-Bouldin (DB) index is

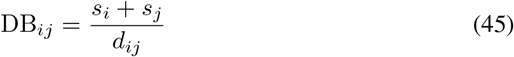

where *s_i_* is the average distance between each point of cluster *C_i_* and its centroid, and *d_ij_* is the distance between the centroids of clusters *C_i_* and *C_j_*. In our experiment, the two clusters are defined as two different distributions of semantic similarities whose centroids correspond to their mean values. The lower DB index is, the more separated the two distributions/clusters are. The lowest possible DB index is zero. Therefore, an DB index close to zero indicates well-separated clusters and thus poorly overlapped distributions. For our experiment, we used the public implementation available in Scikit-learn [67].

Table 8 shows the DB indexes calculated from the results in Figure 12. Here, the low-rank embeddings obtained the best indexes (lowest). Notably, the low-rank embeddings are even able to discover that the groups A and R are weakly related. This is because node pairs labeled as R are indeed weak ancestors (A), as any two nodes in a DAG share a common ancestor at some point, see Figure 12. Finally, it is worth noting that, in comparison to classical techniques especially designed for calculating the semantic similarities between GO nodes (Lin, Resnik, AIC, Wang and GOGO), the Gaussian embeddings are able to obtain much better results, demonstrating the advantages of using the KL divergence between low-rank Gaussian distributions for measuring semantic similarity.

**Table 8:** Davies-Bouldin indexes measuring the level of overlap between the ancestors (A), descendants (D) and rest (R). The two best values are first boldfaced and then underlined, respectively.

| Method | Davies-Bouldin index (↓) | | | | | |
| --- | --- | --- | --- | --- | --- | --- |
|  | BP |  | CC |  | MF |  |
|  | A vs R | D vs R | A vs R | D vs R | A vs R | D vs R |
| Lin | 1.0 | 1.0 | 1.9 | 1.9 | 1.2 | 1.2 |
| Resnik | 1.3 | 1.3 | 2.6 | 2.6 | 1.4 | 1.4 |
| AIC | <u>0.9</u> | 0.9 | <u>1.2</u> | 1.2 | <u>0.9</u> | 0.9 |
| Wang | 2.0 | 2.0 | 4.8 | 3.0 | 2.0 | 1.6 |
| GOGO | 1.4 | 1.4 | 2.0 | 1.4 | 1.3 | 1.1 |
| AROPE | 1.7 | 1.7 | 2.2 | 2.0 | 1.6 | 1.5 |
| GraRep | 335.4 | 355.4 | 13.6 | 13.4 | 23.5 | 26.6 |
| SVD-anc | 2.3 | 2.4 | 8.7 | 4.5 | 4.6 | 3.4 |
| SVD-des | 1.5 | 1.5 | 1.7 | 1.6 | 1.8 | 1.8 |
| DeepWalk | 401.1 | 548.4 | 12.4 | 16.1 | 26.5 | 25.3 |
| LINE | 96.4 | 27.9 | 8.8 | 162.5 | 9.6 | 19.4 |
| node2vec | 4.2 | 4.1 | 3.6 | 3.1 | 3.4 | 3.2 |
| VERSE | 1.3 | 1.3 | 2.0 | 1.4 | 1.5 | 1.3 |
| onto2vec | 7.7 | 7.8 | 23.8 | 12.3 | 9.4 | 29.0 |
| anc2vec | 2.2 | 2.4 | 25.1 | 6.5 | 4.8 | 3.8 |
| neig2vec | 3.2 | 3.4 | 14.6 | 47.2 | 12.0 | 18.2 |
| Graph2Gauss | **0.8** | 0.7 | **1.1** | 0.8 | **0.7** | 0.7 |
| Spherical | 1.2 | 0.7 | 1.2 | 1.0 | 2.3 | 0.8 |
| Diagonal | 1.1 | 0.6 | 1.2 | 0.7 | 0.9 | 0.6 |
| Rank 1 | 1.0 | <u>0.5</u> | 1.6 | <u>0.5</u> | <u>0.9</u> | 0.4 |
| Rank 2 | 1.0 | **0.4** | 1.5 | **0.4** | <u>0.9</u> | **0.3** |
| Rank 3 | 1.0 | **0.4** | 1.3 | **0.4** | <u>0.9</u> | **0.3** |
| Rank 4 | 1.1 | **0.4** | 1.4 | **0.4** | <u>0.9</u> | **0.3** |

